# Batch-effect correction in single-cell RNA sequencing data using JIVE

**DOI:** 10.1101/2023.10.25.563973

**Authors:** Joseph Hastings, Donghyung Lee, Michael J. O’Connell

## Abstract

In single-cell RNA sequencing (scRNA-seq) data analysis, addressing batch effects — technical artifacts stemming from factors such as varying sequencing technologies, equipment, and capture times — is crucial. These factors cause unwanted variation in the data and often obfuscate the underlying biological signal of interest. The Joint and Individual Variation Explained (JIVE) method can be used to extract shared biological patterns from multi-source sequencing data while adjusting for individual non-biological variations (i.e., batch effect). However, its current implementation is originally designed for bulk sequencing data, making it computationally infeasible for large-scale single-cell sequencing datasets. In this study, we enhance JIVE for large-scale scRNA-seq data by boosting its computational efficiency and tailoring it to the single-cell context. Additionally, we introduce a novel application of JIVE which we use to perform batch-effect correction on multiple scRNA-seq datasets. Our enhanced JIVE method aims to decompose scRNA-seq datasets into a joint structure capturing the true biological variability and individual structures which capture technical variability within each batch. This joint structure is then suitable for use in downstream analyses. We employed four evaluation metrics and benchmarked the results against two other popular tools, Seurat v3 and Harmony, which were developed for this purpose. We found that JIVE performed best in metrics that consider local neighborhoods (kBET and LISI) and in scenarios in which the original data contained distinct differences between batches and cell types.

## Introduction

There have been significant advancements in recent years in single-cell RNA sequencing (scRNA-seq) [16]. Single-cell sequencing provides a higher resolution view of genomic data when compared to bulk RNA sequencing (bulk RNA-seq) and allows for the heterogeneity of different cell populations to be preserved. Bulk RNA-seq measures the average level gene expression in a population of cells [31], while scRNA-seq measures the gene expression for each cell individually [30]. Another prominent feature in scRNA-seq data is the high proportion of zero counts [27]. This is due to both biological and technical reasons. One biological cause of this zero-inflation could be due to a particular cell type having very little or no gene expression. A technical cause could be due to what is referred to as a technical dropout, where a certain gene is expressed but does not get detected by the sequencing technology. Multiple sequencing protocols have been developed to capture this information, with prominent examples including CEL-seq2, Drop-seq, MARS-seq, SCRB-seq, Smart-seq, and Smart-seq2 [34]. Sequencing data consists of a count matrix for a given set of genes obtained from a sample of cells. These counts represent the number of times that a specific gene is detected in a single cell [9]. Each row represents a gene and each column is a cell. The library size for a cell is the sum of all counts across all genes.

Once the count data are obtained, it is typically normalized to help account for any variability caused by sampling effects within the given sequencing protocol [9]. A few examples of normalization methods include counts per million (CPM) normalization, upper quartile (UQ) normalization, and trimmed mean of M values (TMM) normalization [27]. In CPM normalization, each count is divided by its cell’s library size and multiplied by a million. UQ normalization uses a scaling factor proportional to the 75th percentile of the counts for a given cell. TMM normalization seeks to trim away cell counts that exhibit large log fold differences within a cell.

Another common preprocessing step is the integration of multiple data sets that are obtained from different batches. Unwanted technical variation and differences between count data are known as batch effects [33]. These effects can arise due to different sequencing technologies being used or cells being sequenced at different times.

### Batch-Effect Correction Methods

There have been over a dozen methods developed to integrate multiple sources of scRNA-seq data that aim to remove these unwanted batch effects [26]. Each method produces a batch-corrected data set which is then used for downstream analyses. Currently, two commonly used methods to correct for batch effects are Seurat v3 [25] and Harmony. [12].

#### Seurat v3

Seurat v3 is software package developed by the Satija lab which provides a comprehensive set of tools for single-cell data analysis and integration [25]. The Seurat v3 integration method builds upon their previous work by leveraging a new graph-based approach. First, log-normalization is performed on all datasets and expression values are standardized for each gene. A subset of features are selected which exhibit high variance across all datasets. Then an initial dimension reduction method utilizing canonical correlation analysis (CCA) is performed to ensure similarities across datasets are preserved. Canonical correlation vectors (CCV) are then approximated and used to identify K-nearest neighbors (KNN) for each cell within their paired dataset. Mutual nearest neighbors (MNN) are then identified to act as anchors between datasets. These anchors are then filtered, scored, and weighted using the new shared nearest neighbors (SNN) approach, and finally used to perform the batch correction.

#### Harmony

Harmony is an integration method designed to model and eliminate effects of known sources of variation [12]. The method utilizes with an initial low-dimensional representation of the data, such as principal components, and then iterates between two algorithms until convergence is reached. The first algorithm clusters cells from multiple batches but ensures that the diversity of batches within each cluster are maximized (i.e., maximum diversity clustering). The second algorithm then uses a mixture model based approach to perform linear batch correction from a given vector of the known batches. The clustering step assigns soft clusters to cells and the correction step uses these clusters to compute new cell embeddings from the previous iteration.

## JIVE

A reasonable alternative to the above methods would be to use a multi-source dimension reduction method, such as joint and individual variation explained (JIVE) [15]. JIVE decomposes two or more biological datasets into three low-rank approximation components: a joint structure among the datasets, individual structures unique to each distinct dataset, and residual noise. The goal in this context would be to use the individual structure to identify the batch effects and the joint structure to identify the biological effects.

JIVE was originally created for integrating different types of bulk sequencing data. For example, [15] used gene expression and miRNA data from a set of 234 Glioblastoma Multiforme tumor cells in an attempt to identify any joint or individual variation between the data types. JIVE can be expressed in terms of principal component analysis (PCA), a common dimension reduction technique. In PCA, given a dataset *X* of size *p × n*, we perform an eigendecomposition on the variance-covariance matrix Σ_*p×p*_ in the following way:

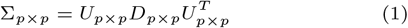

The matrix *U*_*p×p*_ in Equation 1 denotes the *p* eigenvectors (known as loadings) and *D*_*p×p*_ is a diagonal matrix containing the *p* eigenvalues of Σ. We can then calculate a rank *r* approximation of *X* in the following way:

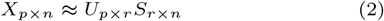

In Equation 2, *U*_*p×r*_ is the first *r* columns from the loading matrix and *S*_*r×n*_ is the first *r* rows from the scores matrix. The *i*-th row of *S*_*r×n*_ is called the *i*-th principal component of *X*, and each principal component is constructed to be orthogonal to all others. Since each principal component is orthogonal, the scores do not suffer from any issues due to multicollinearity. The JIVE decomposition can then be written in the following manner:

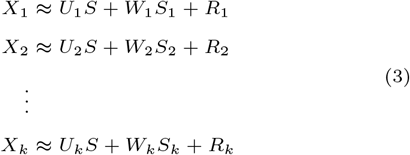

*X*_1_, *X*_2_, …, *X*_*k*_ with *k ≥* 2 denote the original datasets of dimension *p*_*i*_ *× n* where *n* is a common set of objects and *p*_*i*_ is a given set of measurements that can vary for the *k* matrices. We will denote *p*_1_ + *p*_2_ + … + *p*_*k*_ = *p*. Let *r* denote the chosen rank for the equal to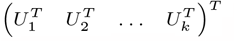, where *U*_i_ are the loadings of the joint structure for the individual datasets *X*_1_, …, *X*_*k*_, and *S* is an *r × n* score matrix. The *W*_*i*_ are *p*_*i*_ *× r*_*i*_ loading matrices and the *S*_*i*_ are *r*_*i*_ *× n* score matrices for each *X*_*i*_. The *R*_*i*_ represent any remaining residual noise. In this paper, we will consider genes as the *n* common set of objects because each cell can be sequenced for the same gene. The JIVE decomposition enforces orthogonality between the joint and individual structures, which ensures that the two structures capture distinct directions of variation within the data.

The JIVE decomposition estimates the joint and individual structures by minimizing the sum of squared error of the residual matrix. Given an initial estimate for the joint structure, it finds the individual structures to minimize the sum of squared error. Then, given the new individual structures, it finds a new estimate for the joint structure which minimizes the sum of squared error. This process is repeated until a given threshold for convergence is reached. The existing implementation allows ranks to be estimated by one of two different methods: a permutation test rank selection and a BIC rank selection [20]. In this paper, we used the permutation test for rank selection.

While JIVE’s current implementation [20] excels at integrating bulk sequencing data from different genomic layers, it is not feasible for integrating large-scale single-cell sequencing datasets, especially those from multiple batches. This incompatibility arises for two main reasons. First, single-cell data often has a substantially larger sample size than previous JIVE applications (i.e., bulk sequencing). Given this magnitude, necessary optimizations are required to mitigate the method’s computational burden. The current R-based implementation also contributes to this computational inefficiency. Second, JIVE was originally constructed for use as a vertical integration method, where different types of omics data that measure the same biological entity are combined (e.g., DNA, RNA, protein, etc.) [6]. In this case, our data shares the same feature space (scRNA-seq), but each batch contains different biological entities. However, JIVE has previously been used for horizontal integration in meta-analysis [10].

To address these challenges, we improved the computational efficiency of the JIVE algorithm by restructuring it using Rcpp and C++. Additionally, we utilized partial singular value decomposition to calculate the leading singular vectors with greater efficiency. Considering that all batches were sequenced over the same gene set, we can leverage JIVE as a horizontal integration method, using the shared variable set to match the batches. We incorporated these modifications within our updated software package. As a result, the new JIVE method can now estimate the joint structure matrix that captures the shared biological structure between scRNA-seq data from different batches and the individual structures that capture technical effects. To assess the performance of the enhanced JIVE for scRNA-seq data batch-effect correction, we benchmarked it against two established methods: Seurat v3 and Harmony using both simulated and real scRNA-seq datasets. The efficacy of each method was evaluated using five distinct batch correction evaluation metrics.

## Methods

### JIVE Algorithm Improvements

The JIVE algorithm was implemented into the R.JIVE R package [20] from the original MATLAB code. The base functions provided in this package can take a substantial amount of runtime to get results (taking upwards of 12+ hours depending on data). We improved the speed of these base functions in two main ways: utilizing partial singular value decomposition in the RSpectra R package [21] and converting frequently used matrix operations into precompiled C++ code using the Rcpp R package [7] and the RcppEigen R package [3] which provides access to the Eigen C++ linear algebra library.

The original R.JIVE code utilizes singular value decompositions (SVD) in many different areas, however only the largest singular values/vectors are used. A full decomposition takes a lot of time and resources to compute and the majority of the output is not used. We switched to using a partial SVD function in the RSpectra R package which returns the largest singular values/vectors of a given matrix. We compared the partial SVD function to the base SVD function that is used in the R.JIVE package. A benchmark was performed on a dataset of size 1000 *×* 1000 generated from a standard normal distribution in which each function call was repeated 100 times. The top 1, 5, and 10 singular values/vectors were computed from each function call.

The other area in which we made significant improvements was in basic matrix operations. We tested two different functions which implemented C++ code to perform matrix multiplication and compared their performance to the default %*% operator in R. One function uses the Armadillo C++ library via the RcppArmadillo R package [8] and the other uses the Eigen C++ library via the RcppEigen R package. The function using RcppEigen allows us to specify the number of CPU cores to utilize when performing computations. We compared the default matrix multiplication operator %*% in R to the implementations in RcppArmadillo and RcppEigen. A benchmark was performed by multiplying two matrices *A* and *B* of size 1000 *×* 1000 generated from a standard normal distribution in which each function call was performed 100 times.

We compared the runtimes of the original R.JIVE functions to the updated versions which implement the changes in 2.1 to assess overall improvements. Two matrices *A* and *B* of size 200 *×* 1000 were generated from a common joint structure matrix, two unique individual structure matrices, and two residual error matrices generated from a standard normal distribution. A visualization of these datasets can be seen in Figure 1.

**Fig. 1.**
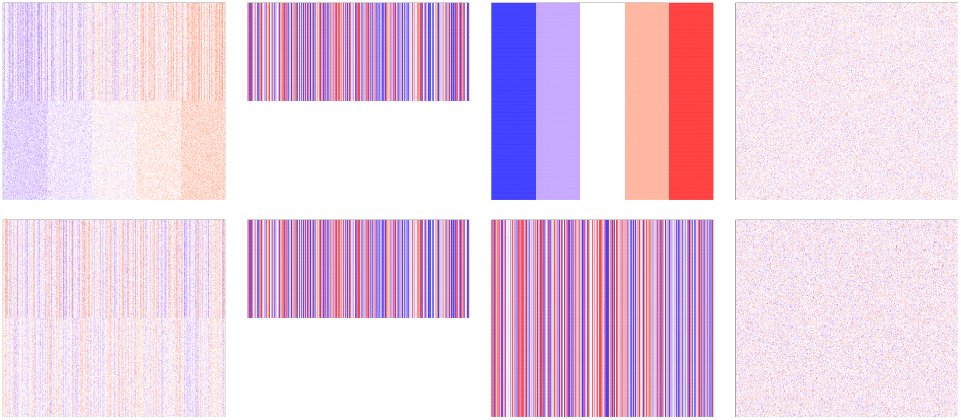
Heatmaps for simulation data used in JIVE benchmarks. The first row shows the final data matrix *A* and the second row shows the final data matrix *B* and their respective decompositions. The first column is the data used as input into the JIVE algorithm. These final datasets were created by adding the second column representing the same joint structure shared between datasets, the third column representing the individual structure unique to each dataset, and the fourth column representing white noise.

The first and second row contain the final matrix in the first column, the joint matrix in the second column, the individual matrix in the third column, and the error matrix in the fourth column for *A* and *B*, respectively. Matrix values range from between approximately -2 (represented by the color blue) and 2 (represented by the color red). A benchmark was performed in which both the old R.JIVE implementation and the updated version were repeated 20 times to record runtimes. We used given ranks of 1 for the joint structure and each individual structure.

All benchmarks were performed on a laptop with a 3.20 GHz AMD Ryzen 7 processor with 16.0 GB RAM using R version 4.2.2.

### scRNA-seq Datasets

One simulated dataset and two real scRNA-seq datasets were used to evaluate the performance of the batch correction methods. The two real data sets were acquired using the scRNAseq R package [23]. Principal variance component analysis (PVCA) [14] was performed for each raw dataset to get an estimation of how much variability is attributed to batch effects, cell type effects, and random error. PVCA uses principal components from PCA and variance components analysis (VCA) to fit a mixed linear model using the factors of interest as random effects to partition the total observed variability into variability due to batches, cell types, or residual error. This helps to quantify the impact of each source of variation on subsequent analyses.

#### Simulated Data

The simulated data was created using the Splatter R package [32] which allows the user to implement a Splat model. The core of the Splat model is a gamma-Poisson distribution which is used to generate a matrix of cell counts for a given number of genes. More than 20 different parameters are available for modifying, including parameters that affect library size, gene means, expression outliers, the presence of batch effects, the size of batch effects, and more. In our dataset, we simulate data for 5000 genes with two batches containing 500 cells each consisting of three different cell types in similar proportions between batches. The frequency of cells in each batch and cell group can be seen in Table 1.

**Table 1.** Batch and Cell Type Frequency for Simulated Data.

| Batch Name | Group 1 | Group 2 | Group 3 | Total |
| --- | --- | --- | --- | --- |
| Batch1 | 312 | 136 | 52 | 500 |
| Batch2 | 294 | 148 | 58 | 500 |

#### Bacher T-Cell Data

The Bacher CD4+ T-cell RNA sequencing data [2] was obtained from six unexposed and fourteen COVID-19 patients. There are a total of fifteen different batches and six different cell clusters provided. This data was chosen as there are not very large distinctions between batches and clusters, so we wished to see how the methods would perform in this type of scenario. The frequency of cells in each batch and cell group can be seen in Table 2.

**Table 2.** Batch and Cell Type Frequency for Bacher T-Cell Data.

| Batch Name | Central Memory | Cycling | Cytotoxic / Th1 | Tfh-like |
| --- | --- | --- | --- | --- |
| 14 | 230 | 1 | 76 | 250 |
| 15 | 239 | 4 | 143 | 425 |
| Batch Name | Transitional Memory | Type-1 IFN signature | Total |  |
| 14 | 190 | 7 | 754 |  |
| 15 | 289 | 13 | 1113 |  |

#### Zilionis Mouse Lung Data

The Zilionis mouse lung data [35] analyzed tumor-infiltrating myeloid cells in mouse lung cancers. There are a total of three different batches and seven different cell clusters provided. This data was chosen as there is some distinct separation due to a batch effect and the cell clusters are well separated. The frequency of cells in each batch and cell group can be seen in Table 3.

**Table 3.**
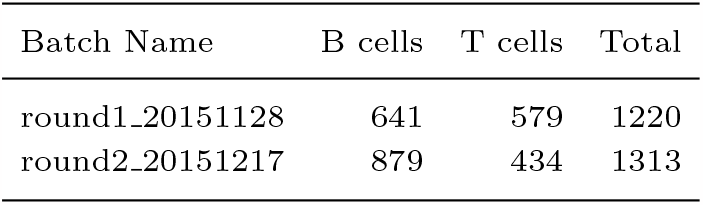
Batch and Cell Type Frequency for Zilionis Mouse Lung Data.

In this study, we evaluate the performance of the three batch-effect correction methods above across three scRNA-seq datasets with four different evaluation metrics. We also present multiple improvements to the JIVE computation algorithms which help significantly decrease runtime compared to the current implementation in the R.JIVE R package.

### Batch Correction Evaluation Metrics

We employed five tools/metrics to evaluate the performance of each of the batch correction methods: visual inspection of t-distributed stochastic neighbor embedding (t-SNE) [28] and uniform manifold approximation and projection (UMAP) [19] dimension reduction plots, k-nearest neighbor batch effect tests (kBET) [4], average silhouette width (ASW) [24], and local inverse Simpson’s index (LISI) [12]. For the visual inspections, we expect to see cells from different batches overlapping each other in the plots with distinct cell type clusters. This is indicative of well-mixed (i.e., integrated) batches that preserve cell type heterogeneity.

While dimension reduction plots are a popular method for evaluating the performance of scRNA-seq batch-effect correction, any conclusions made from them are subjective. We include three numeric evaluation metrics to provide an objective sense of the performance for the three methods.

#### t-Distributed Stochastic Neighbor Embedding

t-SNE is a non-linear dimension reduction technique [28] that aids in visualizing high-dimensional data by assigning each data point a location in a two or three-dimensional map. It aims to preserve as much of the local structure of the original data as possible while also revealing global structure such as clusters. High dimensional Euclidean distances between points are used to create conditional probabilities of one point picking the other as its neighbor. A similar conditional probability is calculated for a low dimensional representation of the data. The goal of t-SNE is to find a low dimensional (i.e., two or three dimensions) representation that matches the two probabilities as best as possible by minimizing a certain objective function. We performed t-SNE using the scater R package [18] version 1.26.1 and the Seurat R package [25] version 4.3.0.

#### Uniform Manifold Approximation and Projection

UMAP is a non-linear dimension reduction technique [19] that is based in manifold theory and topological data analysis. It can be separated into two main phases: graph construction and graph layout. In the graph construction phase, a weighted k-nearest neighbor graph is created, transformations are applied to the graph’s edges, and asymmetry is dealt with. In short, it ensures that the underlying geometric structure of the data is captured. In the graph layout phase, an objective function is defined that preserves important characteristics present in the k-nearest neighbor graph, and the final UMAP representation is the one which minimizes this function. We performed UMAP using the scater R package [18] version 1.26.1 and the Seurat R package [25] version 4.3.0.

#### k-Nearest Neighbor Batch Effect Test

The kBET metric [4] was constructed with the following premise in mind: a subset of a well-mixed dataset with batch-effects removed should have the same distribution of batch labels as the full dataset. A *χ*^2^ -based test is performed for random subsets of a fixed size neighborhood and results from each test (i.e., reject or fail to reject) is averaged over to provide an overall rejection rate. If the rejection rates are low, then we failed to reject most of the tests, and thus the distribution of batch labels in the small neighborhoods were not significantly different from the entire data’s distribution of batch labels. We performed kBET using the kBET R package [5] version 0.99.6. We calculate the rejection rates with neighborhood sizes equal to 5%, 10%, 15%, 20%, and 25% of the number of cells in each dataset. We then use the first 30 principal components from the batch-effect corrected datasets to perform the kBET at each neighborhood size. We then calculate the acceptance rate (1 - rejection rate) so that larger values are more desirable. The acceptance rates are then used for comparison across all methods.

#### Average Silhouette Width

A silhouette is a measure of consistency within clusters of a given dataset [24]. For each data point in a given cluster, we calculate the mean distance between itself and all other points within the same cluster. We also calculate the smallest mean distance between itself and any other data point not in the same cluster. Then a silhouette is the difference of these two values scaled by the largest of the two. A silhouette takes on values between -1 and 1, with values close to 1 indicating that a particular point is appropriately clustered and values close to -1 indicating the opposite. The ASW is the average of all silhouette values which gives a measure of how well-clustered the data are as a whole. We performed ASW calculations using the cluster R package [17] version 2.1.4.

For our purposes, we use the Euclidean distance metric for all calculations. We then subsample our data down to 80% of the original and use the first 30 principal components from the subsampled batch-effect corrected datasets. We calculate two ASW metrics: ASW batch (the batch labels are the clusters) and ASW cell type (cell type labels are the clusters), and this process is repeated 20 times for each method. ASW batch and ASW cell type results from all methods are separately scaled to be between 0 and 1. We report 1 - ASW batch values so that large values are more desirable. The median values of each of these scores are then used for comparison across all methods.

#### Local Inverse Simpson’s Index

The local inverse Simpon’s index [12] first builds local Gaussian kernel-based distributions of neighborhoods around each cell. These neighborhoods are then used in conjunction with the inverse Simpon’s index to calculate a diversity score which corresponds to the effective number of clusters in a particular cell’s neighborhood. We performed LISI calculations using the lisi R package [11] version 1.0.

We calculate two LISI metrics: LISI for batch label clusters (iLISI batch) and LISI for cell type clusters (cLISI cell type). Both LISI scores are calculated for each cell in the batch-effect corrected datasets for each method. iLISI and cLISI results from all methods are separately scaled to be between 0 and 1. We report 1 - cLISI cell type so that large values are more desirable. The median values of each of these scores are then used for comparison across all methods.

### Software and Data Availability

The JIVE implementation used for this analysis can be found at https://github.com/oconnell-statistics-lab/scJIVE. The full code used to perform this analysis can be found at https://github.com/jwhastings/scProject/tree/main. This includes code to replicate the simulated data and to read and process the Bacher T-cell data and the Zilionis mouse lung data.

## Results

### JIVE Runtime Improvements

The runtimes comparing the new implementations of partial SVD and matrix multiplication can be seen in Figure 2. We see that the original functions perform the decompositions in approximately 1.67 seconds on average, while partial SVD functions are performed in 0.08, 0.12, and 0.15 seconds on average. The implementation of a partial SVD function had significant improvements on the overall runtime of the JIVE algorithm. We observed a 95.4% shorter runtime when estimating one singular value/vector, a 93.1% shorter runtime when estimating five singular values/vectors, and a 91.0% shorter runtime when estimating ten singular values/vectors. This is of particular importance because a partial SVD is computed for the joint structure matrix and each individual structure matrix in every iteration of the JIVE estimation algorithm.

**Fig. 2.**
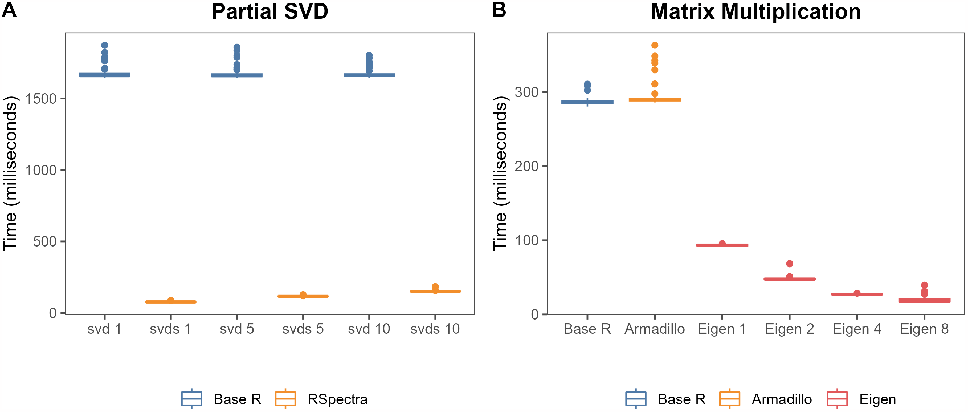
Benchmarks for (A) partial SVD and (B) matrix multiplication runtimes for matrices of size 1000 *×* 1000 colored by their respective R package. The columns in (A) represent the base R function svd estimating *n* singular value(s)/vector(s) and the RSpectra function svds estimating the same, where *n* is one, five, and 10, respectively. The columns in (B) represent matrix multiplication performed using the base R operator %*%, Armadillo via the RcppArmadillo R package, and Eigen via the RcppEigen R package. The last four columns of (B) represent calls to the same Eigen function utilizing one, two, four, and eight CPU cores, respectively.

We can also see the original multiplication operator and the function using RcppArmadillo both averaged just under 0.3 seconds. The next four functions use the RcppEigen utilizing 1, 2, 4, and 8 CPU cores with runtimes of 0.093, 0.047, 0.027, and 0.02 seconds on average, respectively. Performing matrix multiplication using precompiled C++ code also provided a sizable decrease in runtime. The difference between the %*% operator and the function implemented in RcppArmadillo were negligible. The function implemented in RcppEigen not only provided significant improvements over the base R operator, but it also allows for the user to specify the amount of CPU cores to utilize during runtime. We observed 67.5% shorter runtime when using the function with one core, a 83.2% shorter runtime when using two cores, a 90.6% shorter runtime when using four cores, and a 92.9% shorter runtime when using eight cores. Note that using a larger number of available CPU cores does not always provide an increase in speed. Computation on smaller matrices tend to be faster without using multiple cores, but computations on large matrices typically run faster on multiple cores. Overall runtime improvements can be seen in Figure 3.

**Fig. 3.**
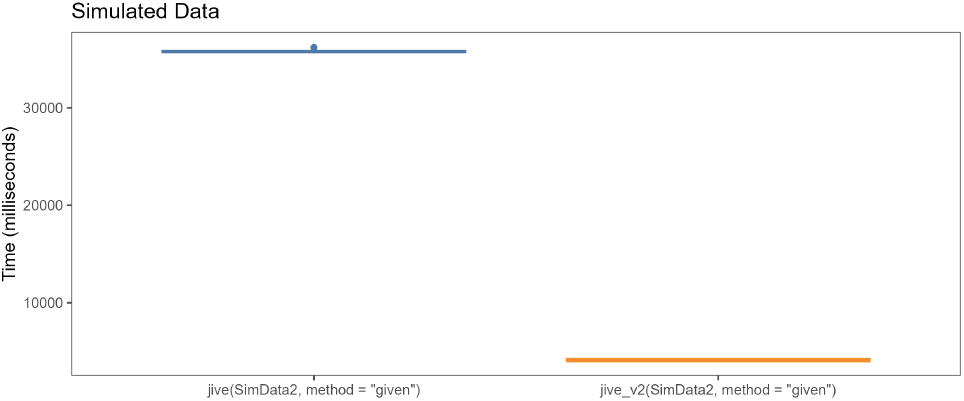
Benchmark for overall JIVE improvements colored by version. The datasets described in 2.1 were used as inputs for the original JIVE function and the updated JIVE function. The first column represents the original JIVE function and the second represents our updated JIVE function.

We see that the original R.JIVE function performs the decomposition in about 35.8 seconds on average, while the improved function completes it in 4.1 seconds on average. The two procedures produced close to identical results. Table 4 shows the proportion of variance attributable to joint structure, individual structure, and residual variance for the two methods.

**Table 4.** Proportion of Variance Attributed to JIVE Decomposition.

| Original | Data 1 | Data 2 | Updated | Data 1 | Data 2 |
| --- | --- | --- | --- | --- | --- |
| Joint | 0.346 | 0.161 | Joint | 0.346 | 0.161 |
| Individual | 0.400 | 0.582 | Individual | 0.400 | 0.582 |
| Residual | 0.254 | 0.256 | Residual | 0.254 | 0.256 |

### Simulated Data

The only preprocessing step performed for the simulated data is a log-normalization [18]. Each cell count is divided by a factor proportional to its library size, a pseudo-count of 1 is added (for zero counts), and a log2-transformation is applied. The PVCA plot for the simulated dataset can be seen in Figure 4.

**Fig. 4.**
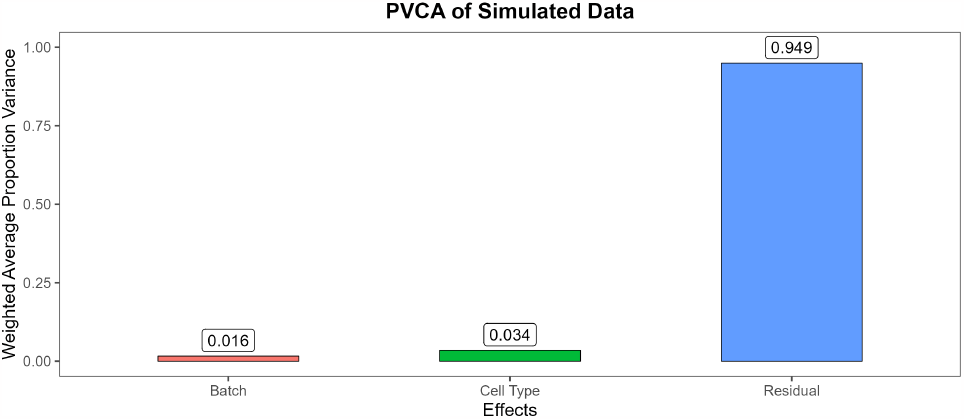
PVCA breakdown for simulated data.

We observe that almost 95% of the variability within the data is not due to the batch or cell clusters which is consistent with most scRNA-seq data. The t-SNE plots can be seen in Figure 5 and the UMAP plots can be seen in Figure 6.

**Fig. 5.**
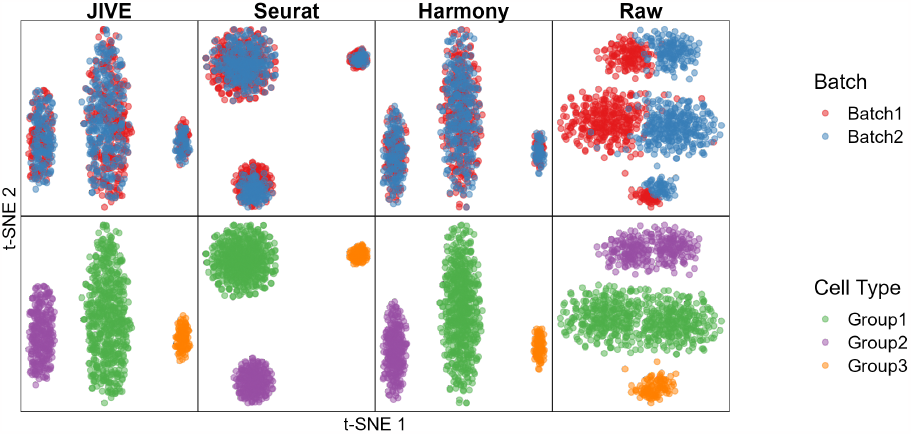
Qualitative evaluation of JIVE, Seurat, and Harmony batch-effect correction methods using t-SNE plots for the simulated data. Each column represents a different method, with the fourth having no batch-effect correction applied. The first row has cells colored by batch and the second row has cells colored by cell type.

**Fig. 6.**
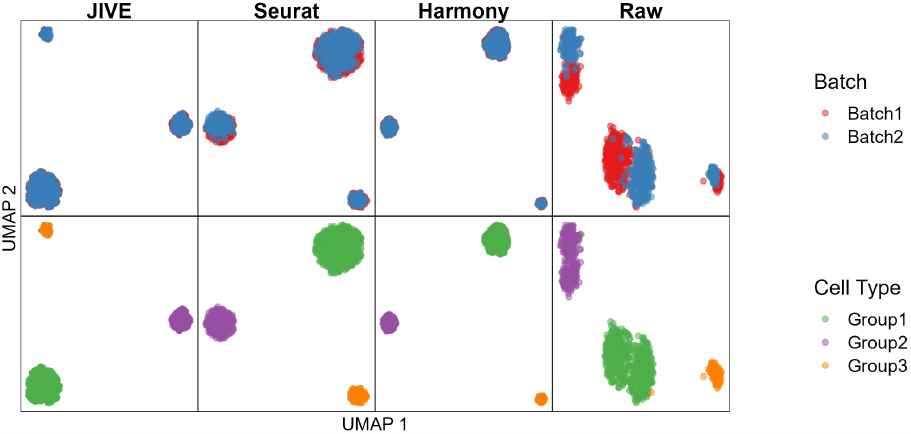
Qualitative evaluation of JIVE, Seurat, and Harmony batch-effect correction methods using UMAP plots for the simulated data. Each column represents a different method, with the fourth having no batch-effect correction applied. The first row has cells colored by batch and the second row has cells colored by cell type.

The top half of each plot has cells colored by batch label and the bottom half has cells colored by their respective cell type/cluster label. Each method has the batches overlapping while the cell clusters are still preserved and distinct from one another. In contrast, the raw plots show some clear separation between batches. The numeric evaluation metrics can be seen in Figure 7.

**Fig. 7.**
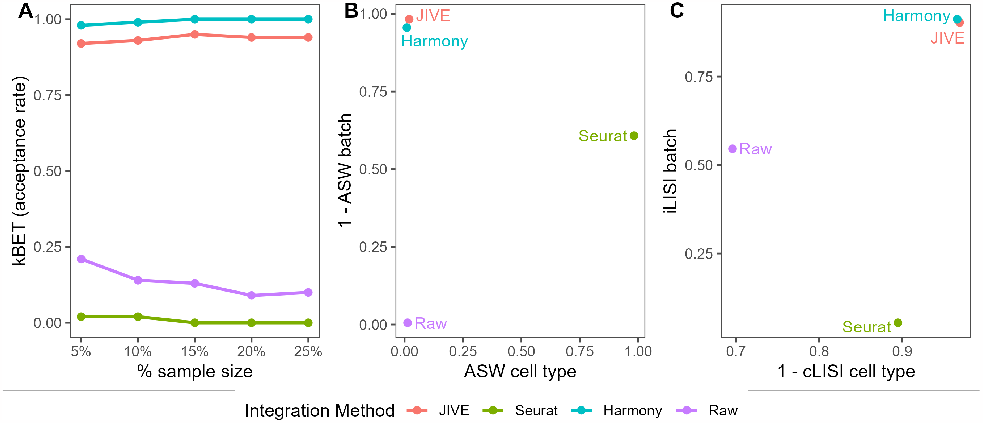
Quantitative evaluation of JIVE, Seurat, and Harmony batch-effect correction methods using (A) kBET, (B) ASW, and (C) LISI for the simulated data. Methods with higher kBET acceptance rates performed best. Methods in the top right of the ASW and LISI plots performed best.

The best performing method for the kBET metric is Harmony followed closely by JIVE. Seurat surprisingly performed worse than the data without any batch correction performed. JIVE and Harmony performed well for the ASW batch metric, closely followed by Seurat. However, Seurat outperformed all methods with regards to the ASW cell type metric. This indicates that Seurat was much better at preserving the different cell types within its cell embeddings than JIVE and Harmony, but not able to distinguish between batches. Harmony and JIVE were the top performers on both the iLISI batch and cLISI cell type metrics. Seurat did not do well with regards to iLISI batch, but was serviceable for the cLISI cell type metric. Overall, JIVE and Harmony were the best performing batch-effect correction methods for the simulated data.

### Assessments Using Real Data

#### Bacher T-Cell Data

Data preprocessing for the Bacher T-cell data consisted of log-normalization (as described for the simulated data), and the top 2000 genes with the highest variability across all batches were selected for analysis. This was performed using a standard workflow suggested in the Seurat R package. We also chose to select only two of the fifteen total batches. This reduced our total dataset size from 33538 *×* 104417 to 2000 *×* 1867. The PVCA plot for the Bacher T-cell dataset can be seen in Figure 8.

**Fig. 8.**
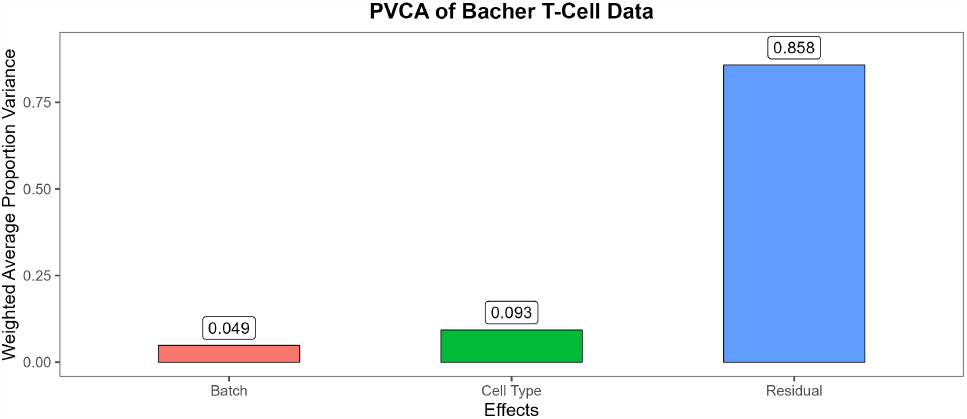
PVCA breakdown for the Bacher T-cell data.

We see more variability attributed to the batch and cell clusters than in our simulated data, with almost 5% and 10%, respectively. The t-SNE plots can be seen in Figure 9 and the UMAP plots can be seen in Figure 10.

**Fig. 9.**
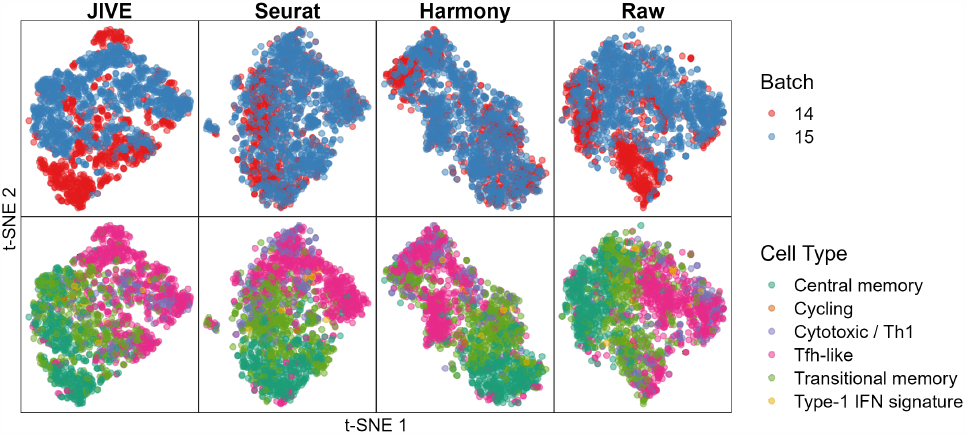
Qualitative evaluation of JIVE, Seurat, and Harmony batch-effect correction methods using t-SNE plots for the Bacher T-cell data. Each column represents a different method, with the fourth having no batch-effect correction applied. The first row has cells colored by batch and the second row has cells colored by cell type.

**Fig. 10.**
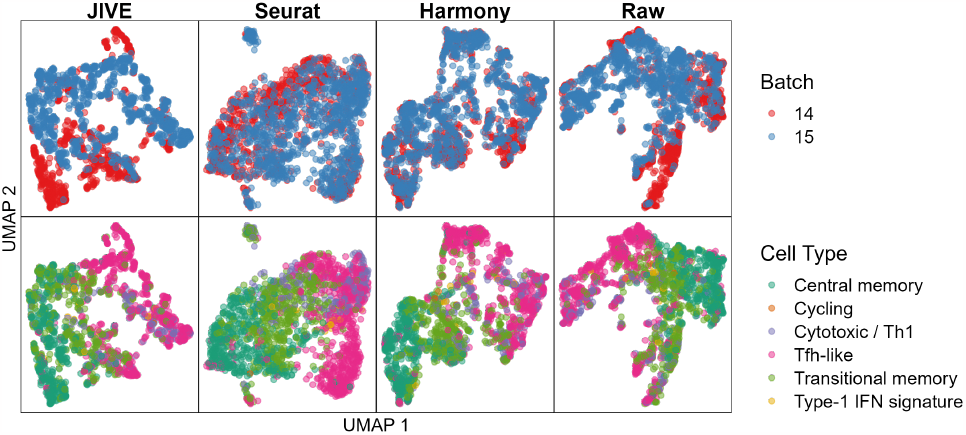
Qualitative evaluation of JIVE, Seurat, and Harmony batch-effect correction methods using UMAP plots for the Bacher T-cell data. Each column represents a different method, with the fourth having no batch-effect correction applied. The first row has cells colored by batch and the second row has cells colored by cell type.

We see that Seurat and Harmony both have well-mixed batches in the dimensionality reduction plots, while JIVE does not. The cell clusters look to be well preserved in all three methods. The numeric evaluation metrics can be seen in Figure 11.

**Fig. 11.**
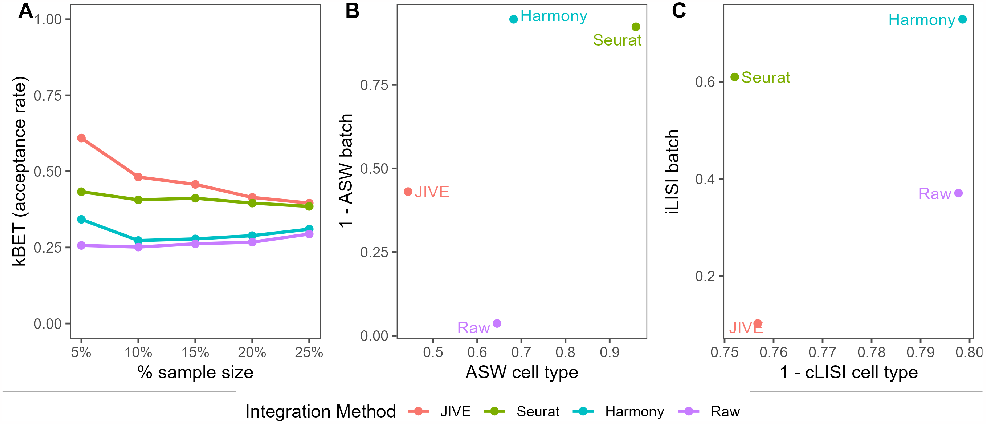
Quantitative evaluation of JIVE, Seurat, and Harmony batch-effect correction methods using (A) kBET, (B) ASW, and (C) LISI for the Bacher T-cell data. Methods with higher kBET acceptance rates performed best. Methods in the top right of the ASW and LISI plots performed best.

JIVE performs the best with regards to kBET, with JIVE close behind. The acceptance rates for Harmony are just slightly larger than the raw data. Harmony and Seurat both perform best in the ASW metrics, while JIVE only outperforms the raw data with regards to ASW batch and actually performs worse in ASW cell type. Harmony is the clear winner in the LISI metrics followed closely by Seurat. Notably, JIVE performs worse than the raw data in both metrics. It is worth noting that the cLISI cell type is on a much tighter scale than the iLISI batch metric in the plot, so the perceived differences are not as large as they appear.

#### Zilionis Mouse Lung Data

We perform the same preprocessing as described for the Bacher T-cell data, select only two batches, and this time selecting only two cell types/clusters (B-cells and T-cells). This helps to reduce the data dimensions from 28205 *×* 17549 to 2000 *×* 2533. The PVCA plot for the Zilionis Mouse Lung dataset can be seen in Figure 12.

**Fig. 12.**
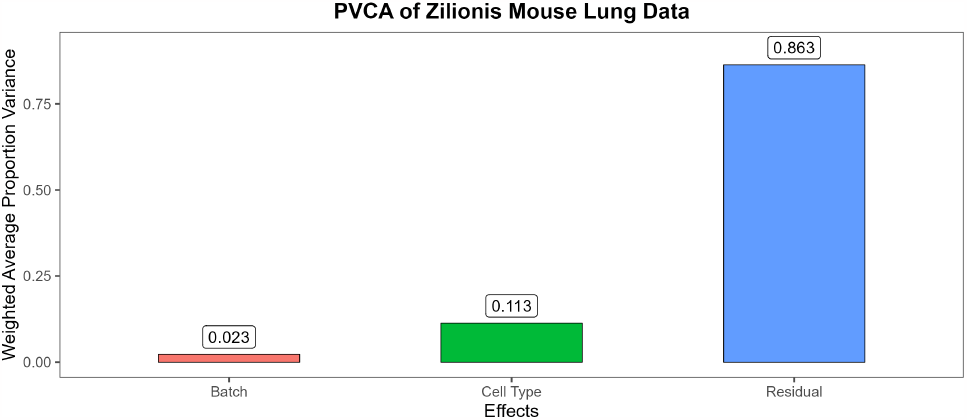
PVCA breakdown for the Zilionis mouse lung data.

We see a similar breakdown of total variability as the Bacher T-cell data except a bit more is attributed to the cell clusters. The t-SNE plots can be seen in Figure 13 and the UMAP plots can be seen in Figure 14.

**Fig. 13.**
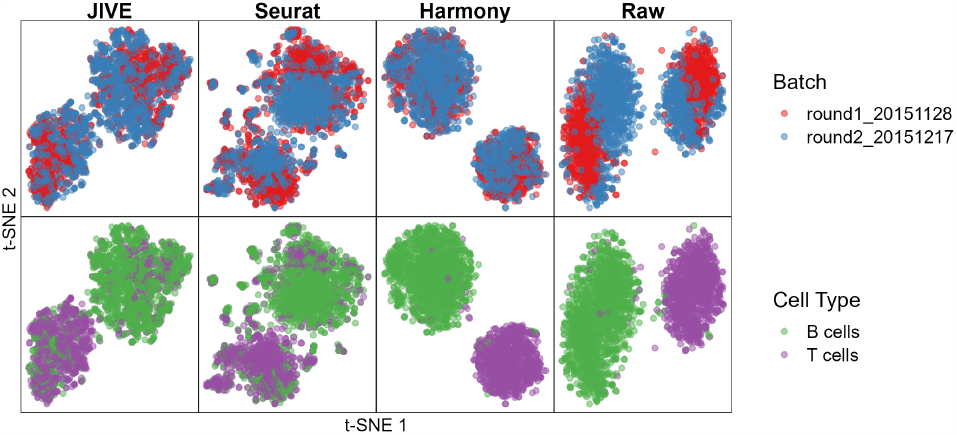
Qualitative evaluation of JIVE, Seurat, and Harmony batch-effect correction methods using t-SNE plots for the Zilionis mouse lung data. Each column represents a different method, with the fourth having no batch-effect correction applied. The first row has cells colored by batch and the second row has cells colored by cell type.

**Fig. 14.**
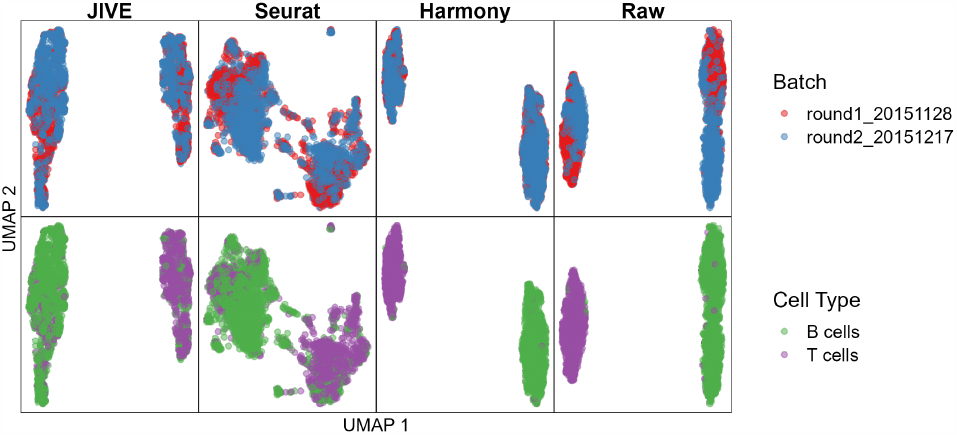
Qualitative evaluation of JIVE, Seurat, and Harmony batch-effect correction methods using UMAP plots for the Zilionis mouse lung data. Each column represents a different method, with the fourth having no batch-effect correction applied. The first row has cells colored by batch and the second row has cells colored by cell type.

We see that each method has well-mixed batches and cell clusters are preserved. It is interesting to note that both JIVE and Seurat produced two distinct clusters that consist of a mix of both cell clusters, while Harmony was able to keep the cell clusters away from each other. The numeric evaluation metrics can be seen in Figure 15.

**Fig. 15.**
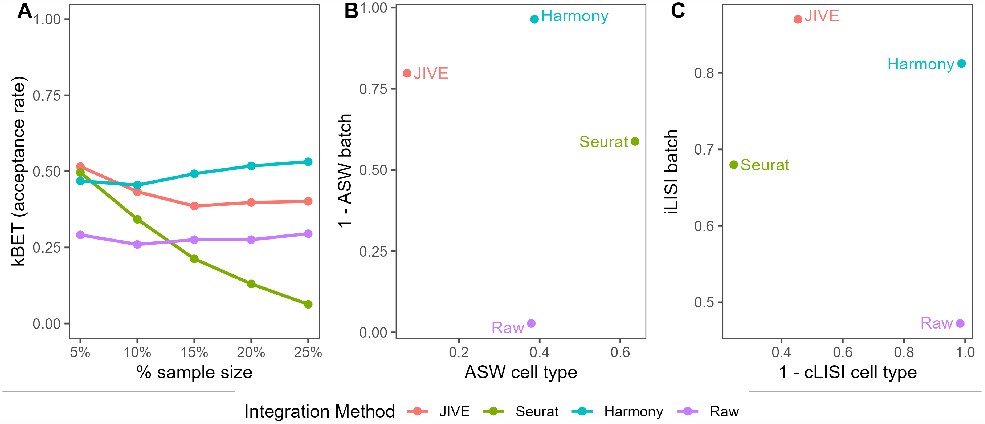
Quantitative evaluation of JIVE, Seurat, and Harmony batch-effect correction methods using (A) kBET, (B) ASW, and (C) LISI for the Zilionis mouse lung data. Methods with higher kBET acceptance rates performed best. Methods in the top right of the ASW and LISI plots performed best.

We see that all three methods perform similarly at low neighborhood levels. However as the size increases, Seurat drops off dramatically. Harmony performs best at kBET with JIVE close behind. The ASW metric performances are a bit of a mixed bag: each method is better than another at either ASW batch or ASW cell type. Harmony performs best at ASW batch, followed by JIVE and then Seurat. Seurat performs best at ASW cell type, followed by Harmony and then JIVE. JIVE and Harmony are the best performers in the LISI metrics, with JIVE being the best with regards to iLISI batch and Harmony winning out in cLISI cell type. Seurat performs worse than both JIVE and Harmony, and only beats the raw data with regards to iLISI batch.

## Discussion

In this paper, we explored if the JIVE method was effective at performing batch-effect correction in scRNA-seq data. Initially, the use of JIVE for this purpose was impractical due to its prohibitively long runtimes with data on the scale commonly seen in scRNA-seq datasets. The significant increase in speed due to the improvements we made to the algorithm make it a much more realistic tool to use for batch-effect correction. We expected JIVE to provide a more flexible alternative to other commonly used methods because not only does it allow the user to specify the ranks chosen for the low-rank approximations, but primarily because it estimates joint structure and individual structure simultaneously. This simultaneous estimation procedure means that JIVE performs both batch correction and dimension reduction at the same time. This is preferred compared to other batch-effect correction methods because less information is lost: any effects not captured in the joint structure will be present in the individual structure, and vice versa. This implies that it is possible to reconstruct the original data using these two estimated structures. This approach differs from Seurat and Harmony where batch correction is performed first and then dimension reduction second. Some information is lost during the batch correction step because only the corrected datasets are estimated and everything else is discarded. One other potential advantage of the simultaneous estimation in JIVE is that one could theoretically use the individual structures as the basis for QC measures to evaluate whether technical effects were truly removed from the joint structure.

There are many other data integration methods which we did not consider in this study. One network-based approach is Similarity Network Fusion [29] (SNF) where multiple sources of data are combined into a single network. It starts by constructing similarity networks for each data source independently and combines them by using a weighted averaging scheme to emphasize that edges are preserved across all networks. It then applies a clustering algorithm to identify groups of nodes that are highly interconnected and likely to be related. A Bayesian-based approach is Multi-Omics Factor Analysis [1] (MOFA) which uses a probabilistic Bayesian and factor anaysis framework to decompose multi-omics data into shared and dataset-specific components. The shared factors are then interpreted in terms of biological processes and their relevance is evaluated.

### JIVE Batch-Effect Correction Performance

In each scRNA-seq dataset, we tested the performance of each batch correction method on their ability to mix batches while still preserving the purity of the cell types/clusters. A commonly used method for evaluating batch integration is by visual inspection of dimension reduction plots, with the most common being PCA, t-SNE, or UMAP plots [26]. This method works well for simple cases like in the simulated data where the batch effects and cell clusters are clearly defined. However, this subjective method tends to become more difficult if there is not clear separation between batches or when cell types are very similar, as is the case in the Bacher T-cell data. This ambiguity that stems from visual inspection is the reason we employed the use of three numeric evaluation metrics to objectively assess the performance of each method. Note that while objective measurements are useful, we still believe that visual inspection can still provide useful insight during exploratory analysis. Note that none of the metrics simultaneously test both the quality of batch mixing and preservation of cell types, and the development of such a metric would be of great interest.

Overall, Harmony performed the best in the simulated data and Zilionis mouse lung data where batch effects were distinct and cell types effects were large. It consistently performed well on kBET and LISI metrics that take into account the structure of each cell’s local neighborhood. The t-SNE and UMAP plots were also consistent with its performance in the numeric metrics. JIVE performed second best with regards to metrics concerning batch mixing, but it struggled with cell type purity metrics. The only dataset where it outperformed the data without batch correction was the cLISI metric in the simulated data. Despite this, it was encouraging to see that JIVE was able to keep up with Harmony in both the simulated data and the Zilionis mouse lung data. Seurat performed second best at metrics concerning cell type purity and had its best performance in the nebulous Bacher T-cell data. The increase in neighborhood size did not have much impact on its kBET performance and it did well in both ASW metrics and both LISI metrics. The most interesting result was Seurat’s poor kBET performance in the simulated data, which seem to contradict the t-SNE and UMAP plots. We recommend using JIVE in scenarios where the original data contains distinct differences between batches and cell types, as it performed best in the simulated data and the Zilionis mouse lung data.

### Limitations and Future Work

Our results showed that there is potential for multi-source dimension reduction techniques to be effective at correcting for batch effects. The major advantage of JIVE over existing methods is that it has a very simple interpretation. However, a major limitation of the JIVE decomposition is that it is best suited to data that is normally distributed. In contrast, scRNA-seq data tends to be zero-inflated and skewed. Therefore, an alternative method that can account for different distributions of data or zero-inflation may be more appropriate. There are some recent methods that can incorporate different distributions. A generalized association study (GAS) is an extension of JIVE that allows for exponential family distributions [13]. A GAS model with a negative binomial link might be a better fit to the scRNA-seq data. Another option would be simultaneous non-Gaussian component analysis, which divides the resulting components into Gaussian and non-Gaussian components [22]. However, more research is needed to develop multi-source dimension reduction methods that can effectively account for zero-inflation.

Another limitation is the narrow scope of datasets used in this study. We only compared three datasets and considered a maximum of two batches at a time with at most six cell types. Subsets of batches in the real datasets were chosen for the sake of simplicity, however in practice it may be of interest to integrate datasets from much more than two experiments or batches. A more comprehensive set of ten different datasets was analyzed in [26], including five distinct scenarios: identical cell types sequenced by different technologies, non-identical cell types across batches, data from more than two batches, big data sets (*>* 100, 000 cells each), and multiple simulated datasets. Assessing the performance of JIVE in a wider range of data scenarios would help give a better sense of its ability to perform batch-effect correction.

## Author contributions statement

D.L. and M.O. conceived the analysis, J.H. conducted the simulations and analysis. D.L., J.H., and M.O. wrote and reviewed the manuscript.

## Acknowledgments

This work was supported by Miami University start-up fund (to D.L. and M.O.) and Shelter Diabetes Research Award (to D.L.).

